# A model for autonomous and non-autonomous effects of the Hippo pathway in *Drosophila*

**DOI:** 10.1101/128041

**Authors:** Jia Gou, Lin Lin, Hans G. Othmer

**Author notes:** The authors declare no conflict of interest. Authors are listed alphabetically. Jia Gou and Lin Lin contributed equally to this work.

## Abstract

While significant progress has been made toward understanding morphogen-mediated patterning in development from both the experimental and the theoretical side, the control of size and shape of tissues and organs is poorly understood. Both involve adjustment of the scale of gene expression to the size of the system, but how growth and patterning are coupled to produce scale invariance and how molecular-level information is translated into organ- and organism-level functioning is one of the most difficult problems in biology. The Hippo pathway, which controls cell proliferation and apoptosis in *Drosophila* and mammalian cells, contains a core kinase mechanism that affects control of the cell cycle and growth. Studies involving over- and under-expression of components in the morphogen and Hippo pathways in *Drosophila* reveal conditions that lead to over- or undergrowth. Herein we develop a mathematical model that incorporates the current understanding of the Hippo signal transduction network and which can explain qualitatively both the observations on whole-disc manipulations and the results arising from mutant clones. We find that a number of non-intuitive experimental results can be explained by subtle changes in the balances between inputs to the Hippo pathway. Since signal transduction and growth control pathways are highly conserved across species, much of what is learned about *Drosophila* applies in higher organisms, and may have direct relevance to tumor dynamics in mammalian systems.

---

The *Drosophila* wing disc is an excellent system for studying the signal transduction and gene control networks involved in growth control, many of which were first discovered there. Growth control in the disc involves both local signals within the disc and system-wide signals such as insulin that coordinate growth across the organism. It is known that the local control network produces correctly-sized discs under various abnormal conditions [1], and we focus on it here. Both disc-wide and clone experiments with various mutants have lead to a rich variety of abnormal growth patterns that remain to be explained in the framework of the known signaling networks, and we will show that these can be understood as the result of subtle alterations in the balances between the outputs of these networks. Since the pathways are tightly linked, the strengths of the interactions determine the outcome, and thus a Boolean on-off description in terms of activation and inhibition of the components is insufficient – a quantitative model is needed.

The Hippo pathway or module is a highly-conserved kinase cascade comprised of the kinases Hippo (Hpo) and Warts (Wts) and the adaptor proteins Salvador (Sav) and Mob as tumor suppressor (Mats) (*cf.* Figure 1). The key effector of this module is Yorkie (Yki) and Wts is its master regulator. Yki is a co-transcription factor whose nuclear access is controlled by Wts via phosphorylation – phosphorylated Yki cannot enter the nucleus. Yki binds to transcription factors such as Scalloped (Sd) to activate the expression of *cyclin E*, *myc*, *DIAP1*, and *bantam*, which control cell proliferation, and it also controls expression of genes upstream of the Hippo module, such as *expanded*, *merlin*, *kibra*, and *four-jointed* ( *fj*) [2]. Two upstream modules provide inputs to the Hippo module, one based on Crumbs, Expanded, Merlin and Kibra that affects the kinase Hippo, and the other based on two atypical cadherins, Fat (Ft) and Dachsous (Ds) that are involved in the regulation of Wts. The former will not be discussed further – it’s input to Hippo is assumed to be constant throughout.

**Figure 1:**
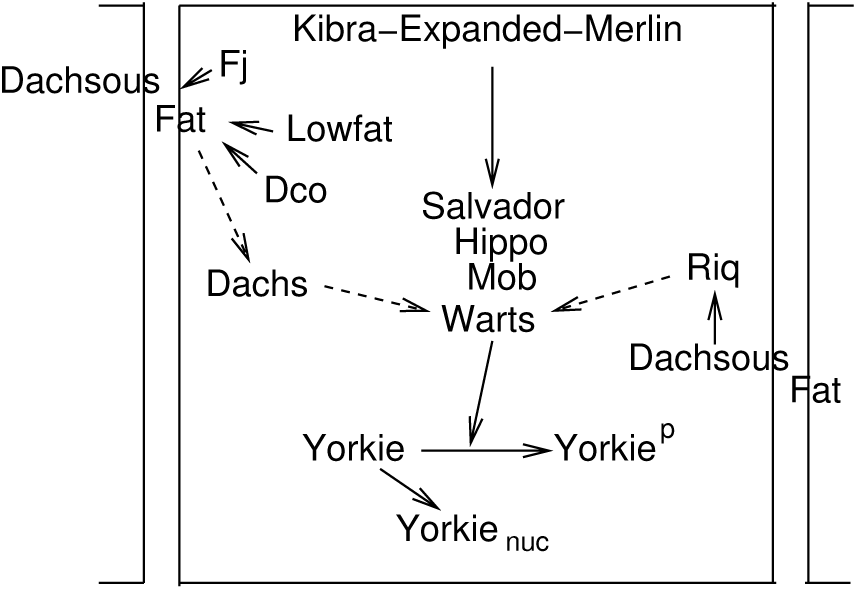
A schematic of the signaling network in contiguous cells. Solid lines denote activation, dashed lines denote inhibition.

Both Ft and Ds are large cadherins with intracellular, transmembrane and extracellular domains. The intracellular domains (ICDs) of each can independently mediate signaling through the Hippo pathway within a cell, while Ft and Ds on adjacent cell membranes can also associate via their extracellular domains (ECDs) to strengthen the signaling and thereby mediate cell-cell interaction. This illustrates a central feature of this system – there are cell autonomous effects controlled by the components in the cytoplasm or nucleus and the ICDs of Ft and Ds, as well as non-autonomous effects caused by binding of an ECD of Ft or Ds to the ECD of its heterophillic partner.

Binding between Ds and Ft is regulated by Fj, which phosphorylates the ECDs of Ft and Ds in the Golgi [3]. Phosphorylation of Ft enhances its affinity to Ds, while phosphorylation of Ds decreases its affinity for Ft [4]. However, the weaker phenotype of *fj* mutants as compared to *ds* mutants – which is discussed later, and the ability of cells expressing high levels of Ft and Ds to associate without Fj, implies that each has a basal affinity for the other [4].

Signaling from the ICD of Ft suppresses growth via Dachs (Dh), an atypical myosin that is epistatic to *fat* in terms of its growth effect. Loss of *dachs* completely suppresses overgrowth or target gene overexpression (OE) induced by the *fat* loss-of-function mutant, and while the mechanism is not clear, one that captures the effect is proposed later. In normal development, Dh is accumulated near the adherens junctions, and localized Dh can bind Wts and promote its degradation [5]. OE of *dachs* increases wing size, while wing size decreases in the *dachs* loss-of-function mutant [6]. In *ds* mutants, strong but nonpolarized membrane localization of Dh is detected, and in *fat* mutants, there is no detectable change in overall Dh protein levels, which indicates that Ft probably affects the membrane localization of Dh. Experiments suggest that while the polarization of Dh controlled by Ft and Ds is essential for planar cell polarity, it is the amount of Dh localized on the membrane that controls cell growth [6]. Since we focus on growth control we ignore polarized expression of Ds and Fj.

Signaling from the ICD of Ds enhances growth by direct interaction with Riquiqui (Riq), a scaffold for protein-protein interactions, and Minibrain (Mnb), a DYRK family kinase [7]. Ds is required for localization of Riq at the apical junctions, and localized Riq potentiates Mnb phosphorylation of Wts, which reduces its activity [7]. While Ds binding to Ft enhances the inhibitory effect of Ft on Dh localization, the complex also increases binding of Riq to Ds and thereby enhances Riq localization. Recent studies suggest that the Ds ICD and Dh also interact [8], and this may be reinforced in the Ft-Ds complex. However, since modulating the expression of either Riq or Mnb does not influence Dh levels or localization [7], it may be that either Ds ICD has independent binding sites for Riq and Dh, or that Ds only interacts with localized Dh.

Experimental results using disc-wide interventions or mutant clones raise several questions concerning how Ft and Ds collaborate to regulate the Hippo pathway. For instance, the effect of Ft on growth is not a strictly decreasing function of the Ft level, as might be expected [9]. OE of *fat* above wild-type (WT) levels decreases the wing size and complete knockout of *fat* increases the size, but a partial knockout of *fat* decreases, rather than increases, the size. Similarly, the effect of Ds is also non-monotonic: loss of Ds results in enlarged wing discs [10], but OE of Ds using Gal4 drivers can either reduce [10, 11] or enhance growth [7]. In addition, double mutants of *fat* and *ds* overgrow more than either of the single mutants [12]. Growth is also non-monotonic in the expression level of *fj*, and when *fj* and *ds* are co-overexpressed, the reduction in wing size is greater than for OE of either separately [10, 11].

Similarly puzzling results emerge when mutant clones are used in a WT disc. Clonal OE of *ds* upregulates Hippo target genes in cells on both sides of the border [13, 11], while *ds* loss-of-function clones upregulate Hippo targets outside, but not inside the clone border [13]. These require both Ft and Dh, since loss of either suppresses the effects. A similar non-autonomous effect arises when Ft is overexpressed [14, 15], but not when it is underexpressed. Further details are given in recent reviews [16, 17, 18, 19, 20, 21, 22], and a summary of experimental observations related to the Hippo pathway is given in Table S3 in the supplemental material (SM).

The Hippo pathway functions as the hub of regulatory mechanisms that control growth and patterning of the wing disc, and therefore, a mechanistic model of it can provide the framework for integrating other pathways. Most current mathematical models of this pathway focus on planar cell polarity [8, 23, 24, 25, 26, 9], while a few touch upon its role in growth [27, 28, 9]. However, none describe the Hippo pathway mechanistically, and thus cannot predict how changes in various components are reflected in cellular growth. Herein we develop a mechanistic model that incorporates both the intracellular interactions of the principal components in the Hippo pathway and the cell-cell interactions via cadherins at the tissue level.

## 1 The Model

The first step in model development and analysis was to determine whether the network in Figure 1 could predict the experimentally-observed outcomes in the absence of the Riquiqui (Riq) effect on Wts. Preliminary qualitative analysis of a model excluding that interaction showed that it could not account for all of the foregoing results, and showed that two additional pathways are essential – one through Ds that activates Yki, and a Ft-independent pathway in which Ds represses Yki. While the biochemical basis for the positive Ds-Riq pathway has been established [7], the existence of the second interaction has not been, but evidence cited above suggests that it is plausible.

Given the complexity of the network, we do not incorporate all species and their interactions in the model, but retain only the central components. These are Ft, Ds, Dh, Riq, Wts, and Yki, which are produced constitutively, and complexes between them. We ignore feedback from Yki to Fj, and fix the total amount of Fj in all forms – thus the model is strictly feed-forward. The signaling network in each cell is shown in Figure 2. Although there are only 6 primary species, many additional species arise as complexes. All species in the model are listed in Table S1, and the reactions and the equations governing their evolution are given in the SM. We model a 1D line of cells and incorporate the processes shown, as appropriate, for each species (*cf.* Figure 3). For example, the Ft reactions in the *i^th^* cell are production and decay in the cytosol, binding to the membranes and binding to Ds on either of the adjacent membranes. A brief summary of the important assumptions underlying in the model is given next – a more detailed description of the model and the experimental justification of the assumptions is given in the SM.

- The ECDs and ICDs of Ft and Ds are phosphorylated at several sites, which affects their activity differently. In the model, the phosphorylation of Ft and Ds catalyzed by Fj refers to the ECD, and this modulates the binding between them. Phosphorylation of the ICDs of Ft and Ds is induced by heterodimer formation, which increases their signaling [10, 29]. We assume that phosphorylation of the ICDs is fast, which implies that the concentration of the phosphorylated form is proportional to the total concentration of each species.
- The inhibitory effect of Ft on membrane-localization of Dh is modeled by a reduction in the Dh binding rate, and is represented by a decreasing Hill function of Ft and all its complexes. Also, the fact that overgrowth in *fat* and *ds* double mutants exceeds that of either single mutant indicates a Ft-independent negative regulatory effect of Ds on growth [30]. Binding of Dh to the ICD of Ds is observed in *Drosophila* scutellum cells [8], and taken together these facts lead to the hypothesis that localized Dh-Ds complexes decay faster than uncomplexed Dh. This hypothesis ascribes a negative regulatory effect of Ds on growth via degradation of Dh in the complex.
- Since Ds is required to recruit Riq to apical junctions, and this is enhanced by Ft-Ds [9], we assume that cytosolic Riq binds directly to either Ds or Ft-Ds on the membrane.

**Figure 2:**
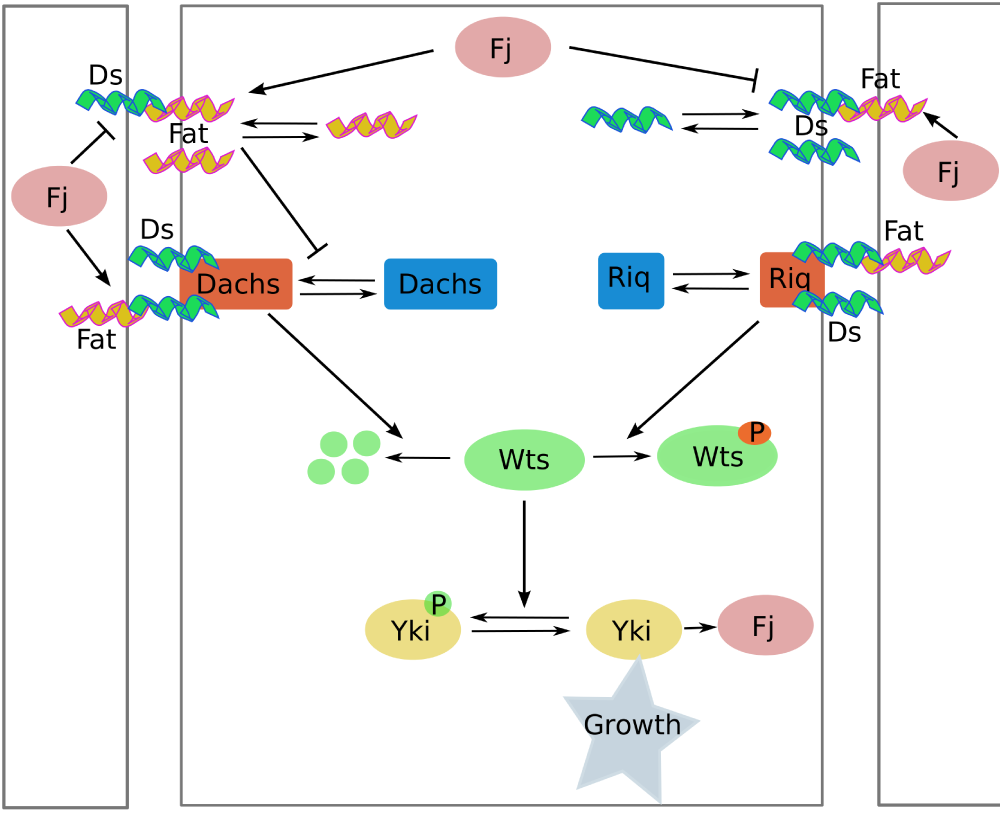
The model signaling network. There is a top-level Ft & Ds module, an intermediate Dh & Riq module, and a terminal Wts & Yki module. The Ft-Dh path depresses Yki, while the Riq-Wts pathway enhances the Yki effect on growth.

**Figure 3:**
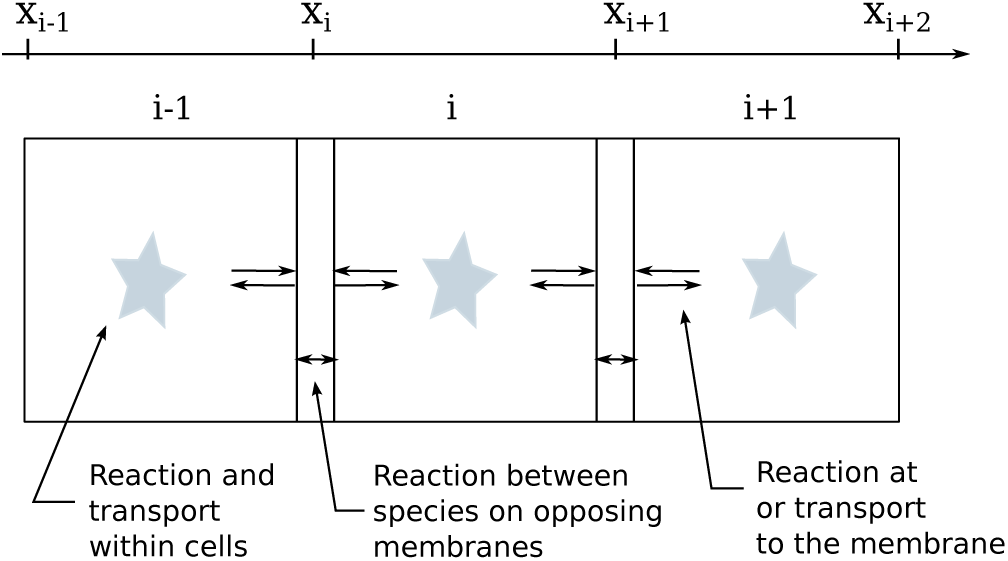
A schematic of a 1D network of coupled cells showing the processes within and between cells. Equations and details are given in the SM.

## 2 Computational Results

The kinetic parameters in the Hippo pathway are currently unknown, and we therefore studied wide ranges of them to understand the properties of the system. We found that the model is insensitive to variation of most of the parameters over 3 orders of magnitude, and most of the observations can be replicated in these parameter ranges. The model parameters are listed in Table S2, and a sensitivity analysis to identify key parameters is discussed later.

### 2.1 The non-monotonic response of Yki in whole discs

Since we assume that Ds and Fj expression is spatially-uniform in WT discs, all cells in the disc, except those at the boundaries, behave similarly. Thus the interactions can be understood by analyzing the signaling network in a single cell in which the reciprocal binding of Ft and Ds between cells is incorporated by identifying the two sides of the cell. This reduction provides a tractable way to explore the behaviors of disc-wide behaviors, including mutants.

Under this assumption the model reduces to a small system of reaction-diffusion equations with nonlinear boundary conditions that is solved for the steady-state concentrations of all species. The predicted Yki concentration as a function of the Ft and Ds production rates is shown in the ’heat-map’ in Figure 4. Although details of this map depend on parameters, it has several important features. Firstly, in *ds^−/−^* mutants the Yki concentration decreases monotonically with the Ft production rate, since the stimulative effect of the Ds-Riq path is absent. In *ft^−/−^* mutants the inhibitory effect of Ft is absent and Yki is regulated by the Ds-Riq pathway and the Dh-Ds interaction. By comparing Figure 5(b) and (a) one sees that overgrowth in a *ds* mutant is slightly weaker than in a *ft* mutant, as is observed experimentally [22]. Another prediction is that double mutants of *fat* and *ds* – (0,0) in Figure 4 – overgrow slightly more than either single mutant, which is also consistent with experimental results. Previous work [12] has shown that knockout of *ds* potentiates the overgrowth in *ft* mutants, but failed to uncover a mechanistic basis. Our explanation is that with or without Ft, membrane-localized Dh is degraded more rapidly when bound to Ds, and therefore in *ds*^−/−^mutants Wts inhibition by Dh is increased, and Yki increases.

**Figure 4:**
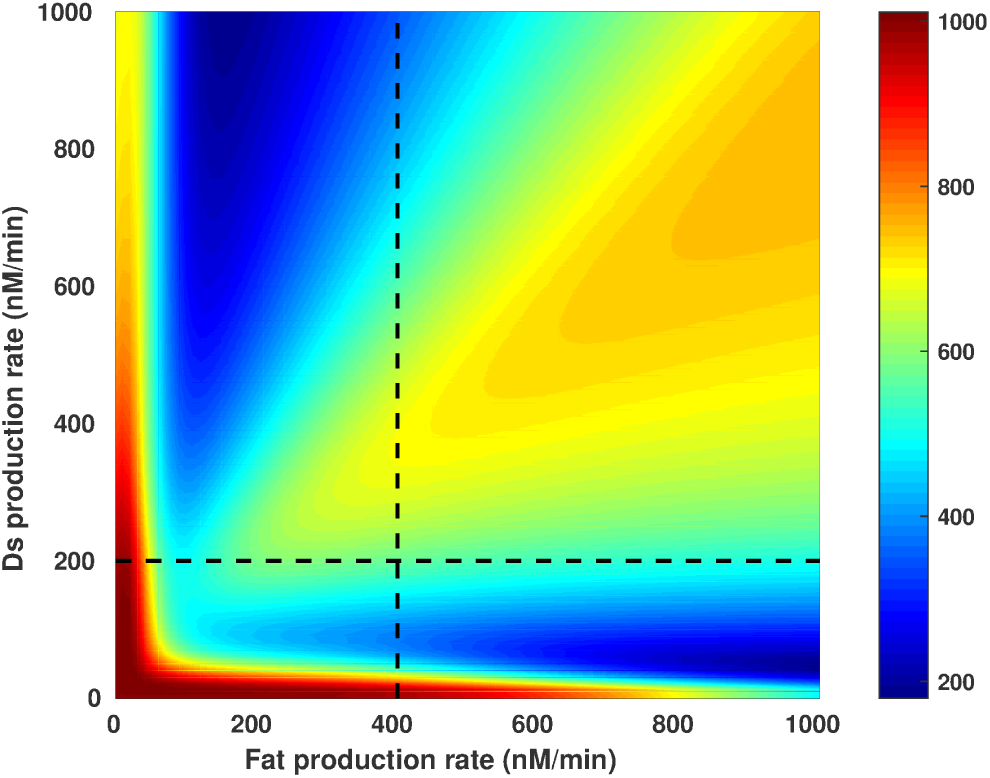
The growth response, as reflected by the Yki concentration, as a function of Ft and Ds production rates. WT rates are (400,200) nM/min.

**Figure 5:**
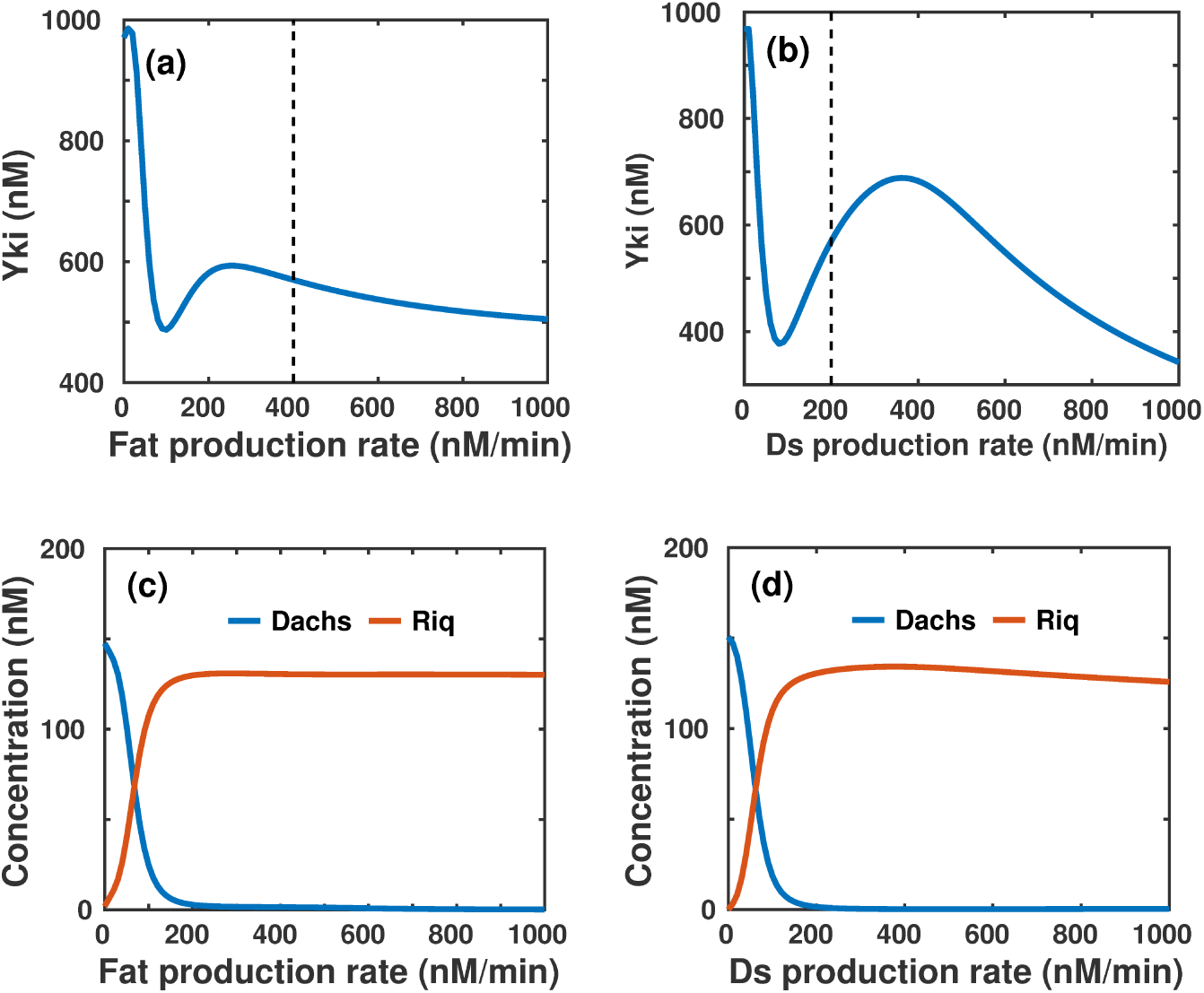
(a) A horizontal slice of the growth response map shows non-monotonic dependence of Yki on Ft expression, and vertical slice (b), shows a similar dependence on Ds expression. (c) & (d) The levels of Wts-bound Dh and its complexes, and Wts-bound Riq complexes, as a function of Ft production (c) and Ds production (d).

In contrast to the monotonic response of Yki in either a Ft or Ds KO, the response of Yki is non-monotonic when Ds expression is fixed at the WT level and Ft production is varied, or conversely, when Ft expression is at WT level and Ds is varied, as shown in Figure 5(a) & (b). Figure 5(a) shows that a Ft KO causes overgrowth, and large OE of Ft causes undergrowth, but in the intermediate range, reduction of Ft production from the WT level first enhances but then reduces, growth. This is in qualitative agreement with experimental observations in [9], where a weak effect on wing size was observed in partial *ft* knockdowns. To understand the non-monotonic response, suppose that the Ft production is increased from zero, and consider the response of Wts bound to Dh and its complexes vs. the response of Wts bound to Riq complexes, as shown in Figure 5(c). The former decreases rapidly due to increased inhibition of Dh localization, which leads to reduced Wts degradation and decreased Yki. On the other hand, the Wts-Riq complexes increase with Ft production, which leads to an increase in Yki. At low Ft the Wts reduction dominates, but at ∼ 100 nM/min the effects balance, and thereafter Yki increases until the level of Wts-Riq complexes saturates at ∼ 220 nM/min, which sets the secondary maximum of Yki. Beyond that the residual level of inhibition via the Ft pathway produces a slow decline in Yki. The balance between the pathways is subtle because Ft affects Dh and Riq through distinct mechanisms and because the inhibitory effects of Ft and Ft-Ds on Dh localization have different strengths.

The model can also explain seemingly contradictory effects of the Ds expression level on growth. Some previous results showed that OE of Ds represses Yki activity [11, 10], but others have argued that it simulates Yki activity [7]. Our results suggest that this disparity may stem from the use of different Gal4 drivers, which may lead to differences in the Ds OE level. A vertical section of Figure 4 at the WT Ft production rate leads to the Yki *vs*. Ds curve shown in Figure 5(b). While strong OE of Ds reduces Yki activity and growth, moderate OE from the WT 200 nM/min to ∼ 400 nM/min increases Yki activity and stimulates growth. Furthermore, the production rate that sets the mid-range maximum growth depends on the Ft production. The model also predicts a non-monotonic effect on growth below WT levels of production. Complete loss of Ds causes overgrowth, and partial loss of Ds reduces growth. This is remarkably similar to the observation when Ft function is lost, emphasizing the similarity of the effects of the two atypical cadherins. These predicted effects can easily be tested experimentally.

One also finds that the Yki level is a monotone increasing function of the Dh or Riq production rate. Furthermore, as the expression level of Dh increases, its degradation of Wts is increased at a fixed Ft level, and as a result Yki activity is promoted. A similar phenomenon is predicted when Riq is overexpressed, as the regulatory effect from the Riq-Ds pathway is positive.

Another interesting phenomenon in the wing disc is cell competition, in which more cells that are more fit by some measure out-compete less fit cells, and proliferate to compensate for the lost cells. This is similar to apoptosis-induced compensatory proliferation [31]. To illustrate what the model predicts we consider a line of 11 cells, five on either side of one that is ”dead” in the sense that all membrane-mediated interactions with neighboring cells are removed and its reactions are stopped. Figure 6 shows the averaged Yki concentration in each cell of the array, and as seen there, the two cells adjacent to the dead cell have a higher Yki concentration and hence would overgrow. This results from two competing effects, the reduction of the inhibitory effect of the Ft-Dh pathway in the WT neighbors due to the loss of Ft*_WT_*-Ds*_D_* binding, and the reduction of the Riq effect due to the loss of Ft*_D_*-Ds*_WT_* binding. Here the former dominates. Qualitatively this prediction agrees well with the cell competition phenomenon observed in experiments.

**Figure 6:**
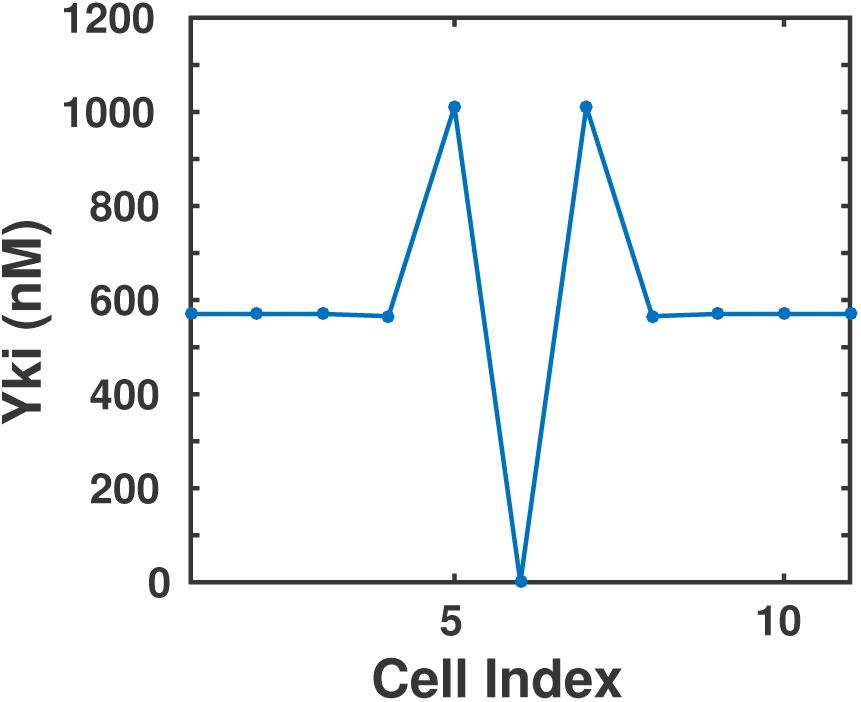
The averaged Yki concentration in an array of 11 cells with one dead cell in the center.

Since the parameters in the Hippo pathway are largely unknown, we investigated how all the foregoing predictions depend on their values. In addition to varying parameters over 3 orders of magnitude and finding that results were insensitive to many of them, we used a more sophisticated sensitivity analysis to identify parameters that most affected the Yki concentration in whole discs. The essence of the variance-based sensitivity analysis described in the SM is to assume that all parameters are independently and uniformly distributed within an interval around their WT value and then analyze how the variance of the model outcome can be decomposed into terms attributable separately to each parameter, as well as to the interactions between them. Figure S3 in the SM shows the first and total order indices of parameters, and as shown there, the steady state cytosolic Yki level is primarily affected by its own production or degradation, and its interaction with Wts. The Riq-Ft-Ds complex and localized Dh are also important in regulating Yki, but the effect of Fj on the Yki responses to Fat or Ds changes is negligible.

### 2.2 Non-autonomous responses

Both Ft and Ds signal autonomously through the Hippo pathway via their ICDs, but they also modulate cell-cell interactions via their ECDs, and we focus on the latter next. Non-autonomous responses – phenotypes induced in wild-type cells by mutant cells – have been observed in a variety of experiments when Ft/Ds signaling is altered by a mutant clone in a WT disc. For instance, OE of Ds in a clone induces hyperactivity of Yki and OE of target genes on both sides of the interface, and the effect vanishes far from the boundary [13]. Figure 7 shows that the model prediction comports with these observations, and Figure 8 shows how localized Dh and Riq are altered in the clone and the adjacent WT cells. In particular, one sees there that the effect of the clone on these species is localized near the clone-WT boundary.

**Figure 7:**
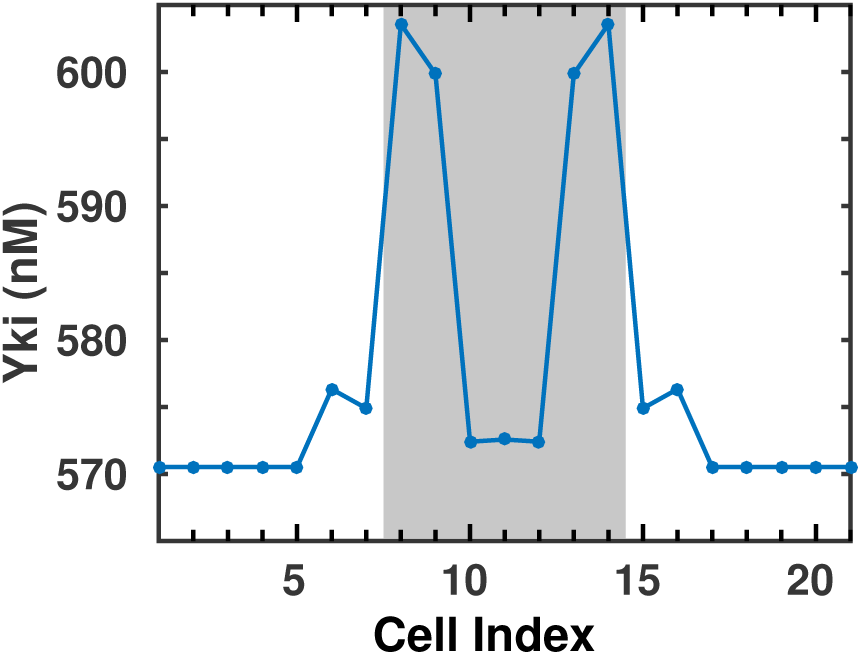
The predicted autonomous and non-autonomous Yki concentration induced by OE of Ds in the clone. A circular array of 21 cells with a patch of 7 clone cells in the shaded region is simulated. The Ds OE level is 2.8xWT.

The elevated Yki level is the outcome of the balance between the inhibitory Ft-Dh pathway and the stimulative Riq pathway. Figure 8(a) shows that the localized Dh level in the two WT cells adjacent to the clone is lower than in WT cells farther from the clone, which reduces its inhibitory effect, while the Riq level is higher in both WT cells. The latter outcompetes and leads to increased Yki.

**Figure 8:**
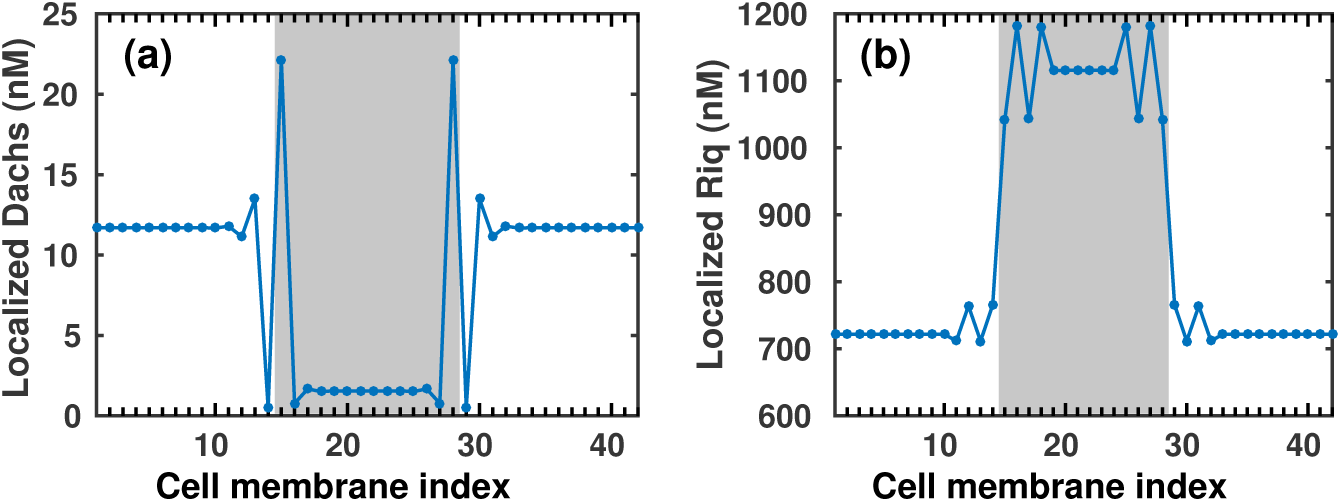
The level of membrane-localized Dh and Riq under Ds OE in a clone. Shown are the 42 locations of cell membranes from 21 cells with 7 clone cells shaded.

In cell 8 of the clone the Dh level on the membrane adjacent to the WT cell 7 is nearly double that of the WT, which suggests strong inhibition of Wts, but the increased localization is in the form of the Ds-Dh complex, which degrades Dh and reduces its inhibitory effect on Wts. Moreover, there is an increase of localized Rig in clone cells that leads to increased Yki levels. Both effects leads to increased Yki.

We also investigated the interaction between WT and clone cells with Ft under expression in the clone to determine how the level of localized Dh at the interface changes in response. As shown in Figure 9, Dh accumulates at the clone boundary due to the reduced inhibition of Ft on Dh binding in the clone, which is expected and as observed in experiments [32]. Furthermore, since we ignore the feedback from Yki to upstream regulators in the model, we investigated the effect of Fj and Dh overexpression in the clone. Increased Dh production induces hyperactivity of Yki as the degradation of Wts is promoted, as shown by the elevated level of Yki in the center of clone cells in Figure 9(b). But as Fj is also overexpressed, the binding of Ft to Ds located on the WT cell membranes is promoted, and the binding of Ft to Ds located on the clone cell membranes is inhibited. Together these account for the increased (decreased) Yki level on the clone (WT) boundary, although the effect in either case is minor.

**Figure 9:**
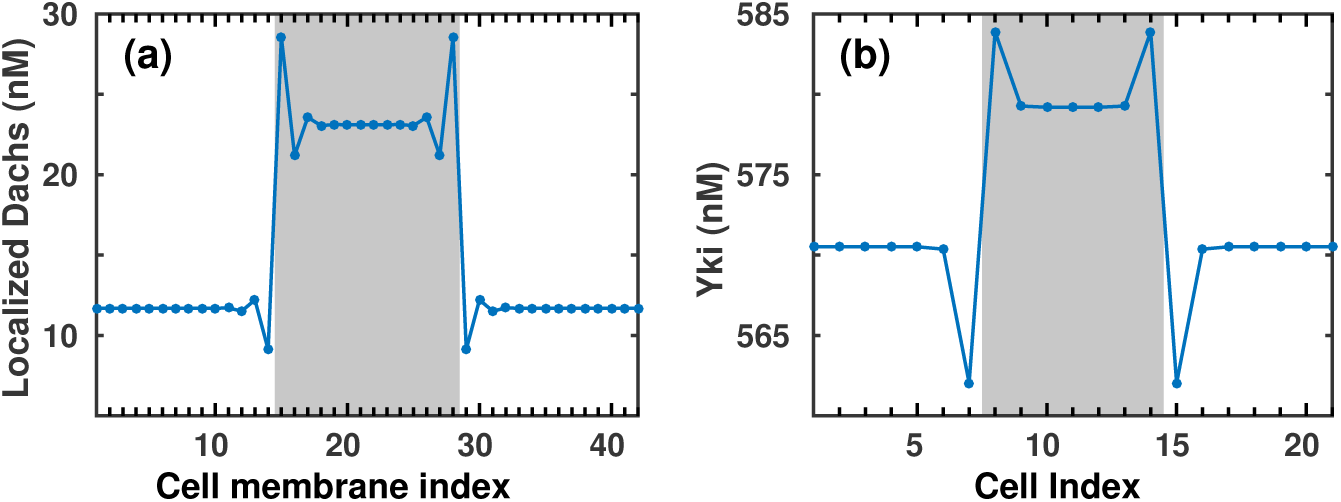
(a) The level of localized Dh with Ft knockdown (0.5xWT) in the clone. (b) The Yki level with Fj at 10xWT and Dh at 2.4xWT in the clone.

Next we consider the case of Ds KO in clone cells. Qualitative analysis of the interactions in the network suggests that a WT cell adjacent to a clone cell will have elevated Yki levels due to the reduced membrane bound Ft*_WT_*-Ds*_C_* complex. In contrast, at the clone side of the interface, the Yki level is suppressed as shown in Figure 10(a). This prediction agrees well with experimental results in which the boundary effect only appears at one side of the interface in Ds knockout clones [13]. Alternatively, since it is not known how insertion of a clone affects the state of the cells at the boundary, we also considered the case in which parameters at the interface change due to interactions between unlike cell types. For instance, in Figure 10(b) the Ft-Ds complex is strengthened at the interface by reducing the off rate of Ft and Ds 5-fold on the adjacent membranes. A comparison with Figure 7 shows that the Yki levels at the two boundary cells are clearly reduced due to the inhibition of Ft-Ds on Dh. Since the nonautonomous responses are caused by cell-cell interactions through the formation of Ft-Ds complexes, we can also explain several other experimental observations. Clearly, in either *fat^−/−^* or *ds^−/−^* mutants, or in double mutants, the non-autonomous responses at the boundary of a clone would disappear, as observed in experiments [13, 15]. Specially, in *ds^−/−^* mutant background, OE of Ds induces the boundary effect only in the Ds expressing cells [13]. Other effects not yet observed experimentally on clone-WT boundary interactions can also be predicted by the model, and these predictions can be tested experimentally.

**Figure 10:**
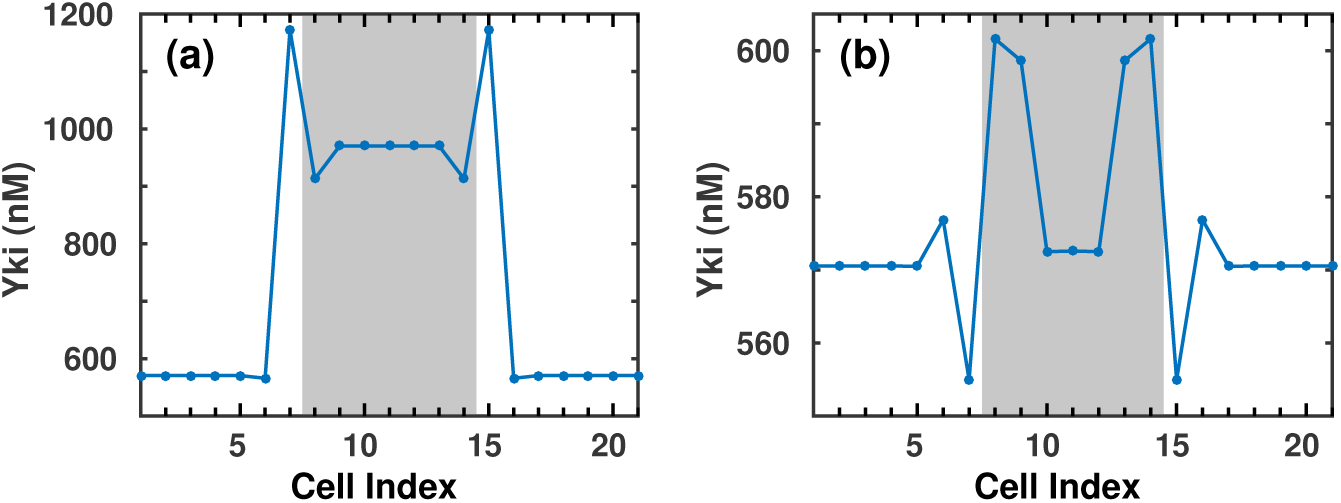
(a) Yki profile in a cell array with Ds knockout in the shaded region. (b) Yki profile in a cell chain with Ds OE in clone cells(shaded region). The unbinding rate between Ft and Ds is reduced at the clone-WT interface.

### 2.3 Boundary signal propagation

The non-autonomous boundary effect due to a clone typically extends ∼2-4 cells away from the clone-WT cell boundary, and in our model, the transmembrane interaction between Ft and Ds controls cell-cell communication. The intracellular processes regulate signal transduction within a cell, and together, the two determine how far the boundary effect propagates. To better understand this we considered a general model – described in the SM – of reciprocal binding processes between two species to explore the impact of parameters on signal propagation. Specifically, we investigated how transport within a cell (diffusion), binding processes (speed and affinity), and degradation processes (decay rates) affect the boundary effect. The model predictions show that the boundary effect induced by OE of Ds in cell clones, including autonomous and nonautonomous responses, are remarkably stable as the nature of signal propagation does not change when the parameters for different kinetic processes are changed within an order of magnitude. It should be noted that in reality signal transport in a cell is probably more complex than simple diffusion, since other processes such as transport along microtubules may be important.

#### Diffusion

Two modes of transport by diffusion are incorporated in the model, one implicitly and one explicitly. Diffusion can occur on the cell membrane, and certainly plays a role in the formation of heterodimers of Ft and Ds. However, since these are bimolecular reactions, the diffusion needed to localize molecules for binding is implicitly incorporated in the binding rate, as is commonly done in such reactions. In the model we only incorporate diffusion in the cytosol, and thus, as expected, with faster diffusion the signal propagates farther from the boundary between wild-type and clone cells, as shown by comparing Figure 11(a) (faster diffusion) and (b). When one side of the cell detects the abnormality of its neighbors via the heterodimer formation process on the membrane, the corresponding change on the other side is more sensitive when diffusion is faster.

**Figure 11:**
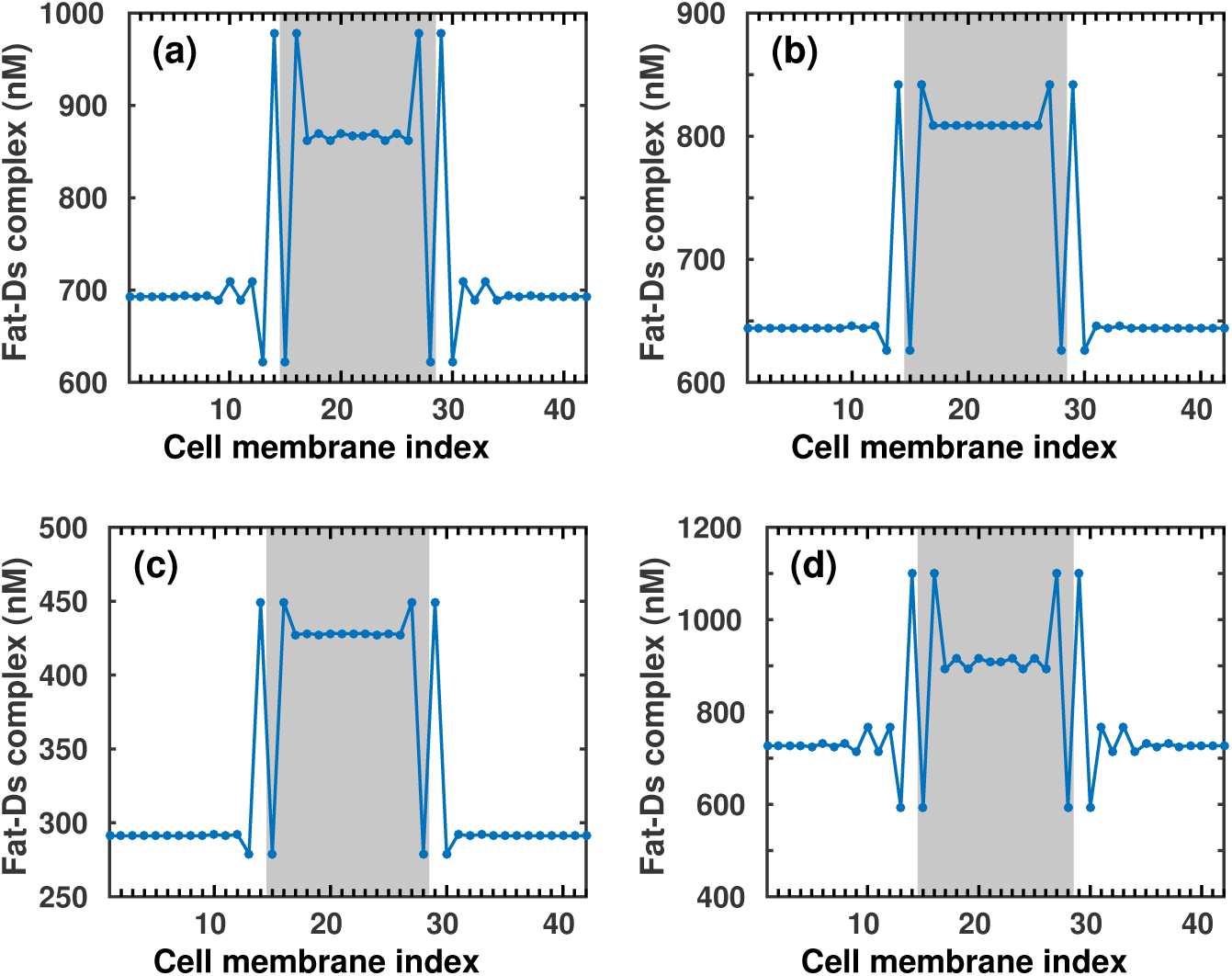
Predicted boundary signal propagation as different kinetic processes change. From (a) to (b): diffusion coefficients of Ft and Ds in the cytosol are decreased from 10 to 1*μm*^2^min^−1^. From (a) to (c): decay rates of Ft and Ds in the cytosol are increased from 1 to 10min^−1^. From (a) to (d): membrane localization processes of Ft and Ds are accelerated by 10 fold. Parameters for the base case (a) can be found in the SM.

#### Degradation processes

We also considered the impact of degradation rates on signal propagation, and find that they affect the boundary effect in a reverse manner, i.e. increasing degradation rates reduce the degree and/or spatial extent of the nonautonomous response, and vice versa. Detailed illustrations of the predicted responses are shown in the SM. Here we show an example of increasing the decay rates of Ft and Ds in the cytosol (Figure 11(c)).

#### Membrane localization

To explore how membrane localization processes influence signal propagation, we examine several cases. Figure 11(d) shows the result of increasing both on (from cytosol to membrane) and off (from membrane to cytosol) rates at a fixed ratio. One sees that expediting membrane localization processes enhances the boundary effect on both sides of the boundary in extent and in amplitude, due to the increase of mass flux from cytosol to membrane. Other simulation results can be found in the SM.

#### Relative concentration

The non-autonomous effect has been reported to be maximal when Ds and Ft are equally expressed [9], which is supported by our simulations. We find that the extent to which the boundary signal propagates, indeed, is maximal if the production rates of Ds and Fat are equal, as shown in Figure 12.

**Figure 12:**
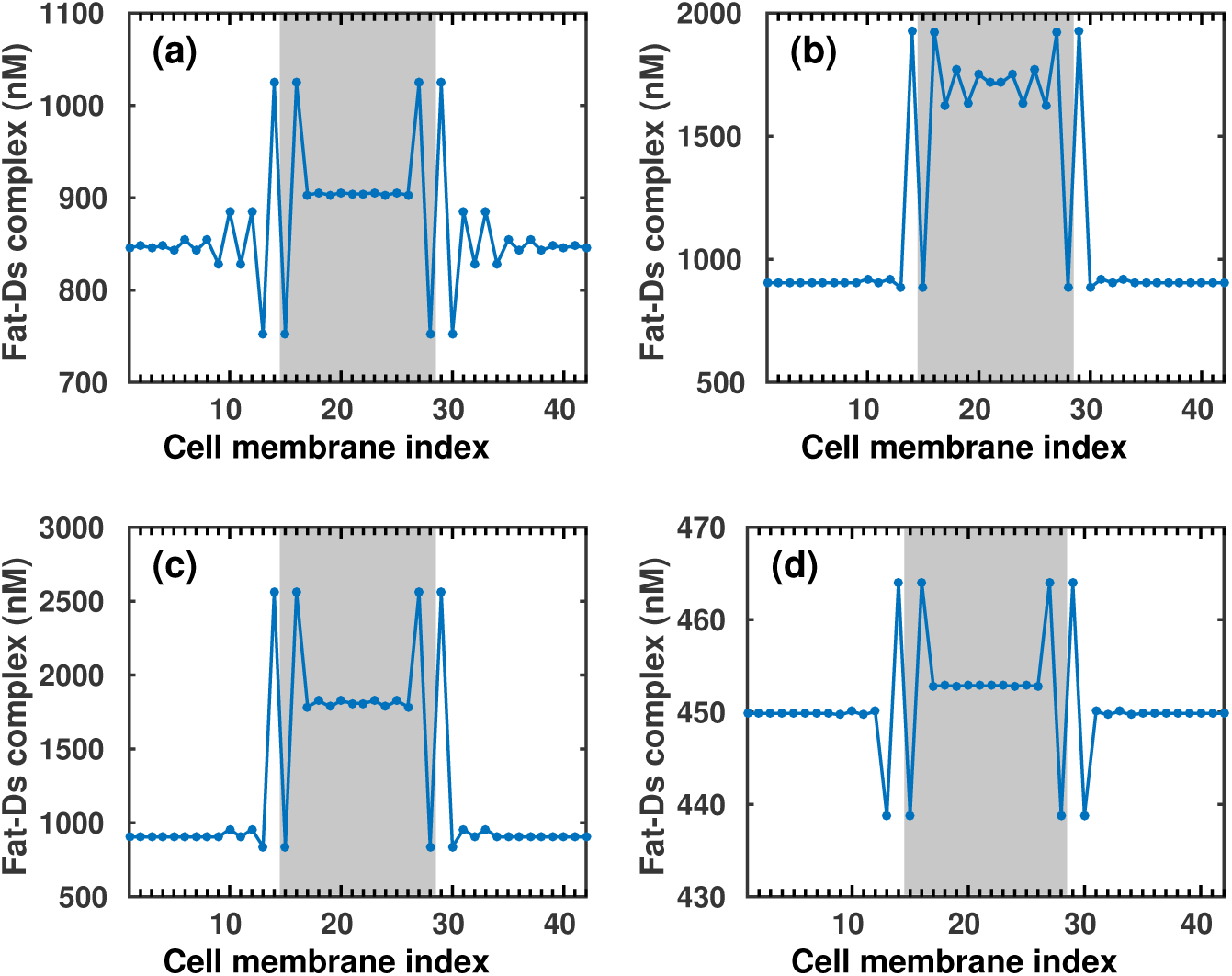
The effect of the relative concentrations of Ft and Ds on boundary signal propagation. (a): Ft and Ds are equal in wild-type cells, and Ds 2xWT in clone cells. (b) Ft is twice of Ds in wild-type cells, and Ds is increased to be equal to Ft in clone cells. (c) Ft is twice of Ds in wild-type cells, and Ds is increased to be higher than Fat in clone cells. (d) Ft is half of Ds in wild-type cells, and Ds doubles in clone cells

## 3 Discussion

The goal of this work was to provide a framework for understanding the complex phenotypes associated with the Hippo pathway and to make testable predictions that can guide further experimental studies. The model developed here can replicate all primary experimental observations, such as the non-monotonic effects in disc-wide alterations of Ft and Ds expression, and the non-autonomous effects induced by cell clones. The model suggests that the seemingly inexplicable observations that result derive from the perturbation of the delicate balance between positive and negative control of intra- and intercellular signals. In particular, we showed that the regulation of Dh and Riq localization on the membrane plays a central role in both non-autonomous and non-monotonic effects (Figure 8). The model also predicts a difference between the autonomous and non-autonomous responses stimulated by clone cells with disrupted Ft/Ds signaling, and provides a mechanistic explanation for the *ft*, *ds* double mutant phenotype, which supports our hypothesis that Ds interacts with Dh. The fact that the model predicts most of, if not all, characteristic phenotypes, demonstrates the applicability of the model to the Hippo pathway.

The non-monotonic response of Ft on growth and the non-autonomous response induced by OE of Ds in cell clones has also been explained by a recent model that assumes mutual inhibition between the opposite orientations of the heterodimers and self-promotion of the same orientations [9]. While this is an interesting hypothesis, there is little experimental evidence in support of it. In contrast, the model developed herein does not assume such roles, and yet predicts both the non-monotonic and nonautonomous responses. These stem from the balances between the positive regulatory step from Ds via Riq, and the interactions between Ds and Dh that is found *in vitro*, but not yet *in vivo.*

While all the major experimental observations can be explained in the one dimensional model developed here, there may be subtleties that will arise in a 2D model, and this will be investigated in future work. Additional factors to be considered are other transport mechanisms, other components in the signaling pathways, and most importantly, the interaction between biochemical signaling and mechanical interactions between cells.

## Acknowledgement

Supported in part by NIH Grant # GM29123.

